# Immunogenicity and In vivo protection of a variant nanoparticle vaccine that confers broad protection against emerging SARS-CoV-2 variants

**DOI:** 10.1101/2021.06.08.447631

**Authors:** James Logue, Robert Johnson, Nita Patel, Bin Zhou, Sonia Maciejewski, Haixia Zhou, Alyse Portnoff, Jing-Hui Tian, Marisa McGrath, Robert Haupt, Stuart Weston, Holly Hammond, Mimi Guebre-Xabier, Carly Dillen, Joyce Plested, Shane Cloney-Clark, Ann M Greene, Mike Massare, Greg Glenn, Gale Smith, Matthew Frieman

## Abstract

The severe acute respiratory syndrome coronavirus-2 (SARS-CoV-2) continues to spread globally. As SARS-CoV-2 has transmitted from person to person, variant viruses have emerged with elevated transmission rates and higher risk of infection for vaccinees. We present data showing that a recombinant prefusion-stabilized Spike (rS) protein based on the B.1.351 sequence (rS-B.1.351) was highly immunogenic in mice and produced neutralizing antibodies against SARS-CoV-2/WA1, B.1.1.7, and B.1.351. Mice vaccinated with our prototype vaccine NVX-CoV2373 (rS-WU1) or rS-B.1.351 alone, in combination, or as a heterologous prime boost, were protected when challenged with live SARS-CoV-2/B.1.1.7 or SARS-CoV-2/B.1.351. Virus titer was reduced to undetectable levels in the lungs post-challenge in all vaccinated mice, and Th1-skewed cellular responses were observed. A strong anamnestic response was demonstrated in baboons boosted with rS-B.1.351 approximately one year after immunization with NVX-CoV2373 (rS-WU1). An rS-B.1.351 vaccine alone or in combination with prototype rS-WU1 induced protective antibody- and cell-mediated responses that were protective against challenge with SARS-CoV-2 variant viruses.

## Introduction

The emergence of severe acute respiratory syndrome coronavirus (SARS-CoV-2) in late 2019 generated an urgent medical need for an effective vaccine to combat the pandemic. SARS-CoV-2 has infected over 138 million individuals causing almost 3 million deaths by mid-2021 (www.who.org). This pandemic has altered economic and social structures throughout the world. Enhanced understanding of coronavirus biology from past outbreaks of SARS-CoV in 2003 and middle east respiratory syndrome coronavirus (MERS-CoV) in 2012 enabled rapid design and emergency use approval of COVID-19 vaccines (*1-5*).

Global spread of SARS-CoV-2 has been punctuated by the identification of mutant viruses, or variants, that have gained function as they spread. As coronaviruses replicate, they have a 1 in 100,000 chance of mutating a base in their genomes due to the high fidelity of the RNA dependent RNA polymerase encoded in the virus (*6, 7*). When selective pressure is placed on the virus, gain-of-function mutations emerge that facilitate improved receptor binding, replicating to higher titers resulting in enhanced transmission, or immune evasion, and the variants that emerge from these bottlenecks have spread with potential to evade current countermeasures. Two variants of concern have emerged in the past year, one first identified in the United Kingdom (UK) called B.1.1.7 and the other in the Republic of South Africa (SA) called B.1.351. B.1.1.7 transmits more readily between individuals than the progenitor strains do, causing increased hospitalizations and prevalence around the world (*8*). B.1.351 has spread to many countries; it rapidly becomes among the most prevalent circulating strains. B.1.1.7 has been shown to be neutralized by SARS-CoV-2 convalescent sera comparably to its progenitor virus, however, B.1.351 has gained the ability to evade several authorized monoclonal antibodies and shows reduced neutralization by convalescent sera *(12,13)*.

A Phase 3 trial in the United Kingdom using our prototype SARS-CoV-2 vaccine, NVX-CoV2373, demonstrated it to be highly effective (86.3%) at preventing infection by the highly transmissible B.1.1.7 strain that was circulating during the trial (*9*). A Phase 2b trial of NVX-CoV2373 in South Africa showed this vaccine’s reduced efficacy against B.1.351 (60.1% in HIV-negative subjects); this decreased efficacy against B.1.351 has also been observed for other SARS-CoV-2 vaccines (*10-13*). In response to these breakthrough transmission events, we developed a full-length, stabilized recombinant spike protein (rS) antigen with the mutations in the B.1.351 variant (rS-B.1.351). We evaluated cellular and humoral immunogenicity and protective efficacy of rS-B.1.351 alone or in combination with our prototype vaccine based on Wuhan-Hu-1 (rS-WU1). B and T cells were highly activated by rS-B.1.351 vaccination leading to the production of neutralizing antibodies and expansion of multifunctional T cells. Immunization with rS-B.1.351 induced high levels of neutralizing antibodies against multiple variant strains of SARS-CoV-2 as well as the parent WA1 strain. Mice immunized with either rS-B.1.351 or rS-WU1, then challenged with B.1.1.7 or B.1.351, showed a 5-log reduction in B.1.351 virus titers and 2-log reduction of B.1.1.7, both to lower than the limit of detection. All immunized mice were protected against challenge-induced weight loss when infected with B.1.351. In olive baboons subjected to a primary immunization series with rS-WU1, boosting with rS-B.1.351 approximately one year later led to a strong anamnestic response, with the rapid induction of anti-Spike IgG and spike-specific T cell responses. Finally, sera from humans vaccinated with NVX-CoV2373 (rS-WU1) were analyzed for their ability to neutralize B.1.1.7 and B.1.351. We found that human sera from trial participants demonstrated neutralizing capacity of B.1.1.7 equal to rS-WU1, with modest reduction in neutralizing capacity against B.1.351.

In this work, we demonstrate the development and evaluation of a B.1.351 Spike directed variant vaccine that induced highly potent neutralizing antibodies, activated both B and T cells, protected mice from clinical disease, and prevented detectable SARS-CoV-2 replication in the lungs. Positive results in non-human primates also suggests the feasibility of utilizing this variant vaccine as a booster after previous immunization with a vaccine directed toward the ancestral SARS-CoV-2 Spike protein.

## Results

### Production of Spike glycoprotein vaccine rS-B.1.351

The antigen component of our prototype vaccine, NVX-CoV2373, was based on the Spike glycoprotein from the SARS-CoV-2/Wuhan1 sequence (now the B.2 lineage; MN908947.3) with mutations: 682-RRAR-685 in the furin cleavage site was mutated to QQAQ to resist proteolytic cleavage (“3Q” mutation), plus K986P and V987P were introduced to increase stability (“2P” mutations). The Spike ORF was cloned into recombinant baculovirus and expressed in Sf9 insect cells as previously described (*14*). NVX-CoV2373 is currently under Phase 3 testing in the US and Mexico, and has completed Phase 3 testing in the UK. Using an identical expression and purification system with the same 3Q-2P mutations, we have also developed a Spike glycoprotein vaccine based on the Spike ORF from the SARS-CoV-2 B.1.351 lineage that emerged in South Africa, also containing the 3Q-2P mutations (Figure 1A). This recombinant spike protein (rS-B.1.351) was expressed, purified, and formulated with Matrix-M1 adjuvant.

**Figure 1.**
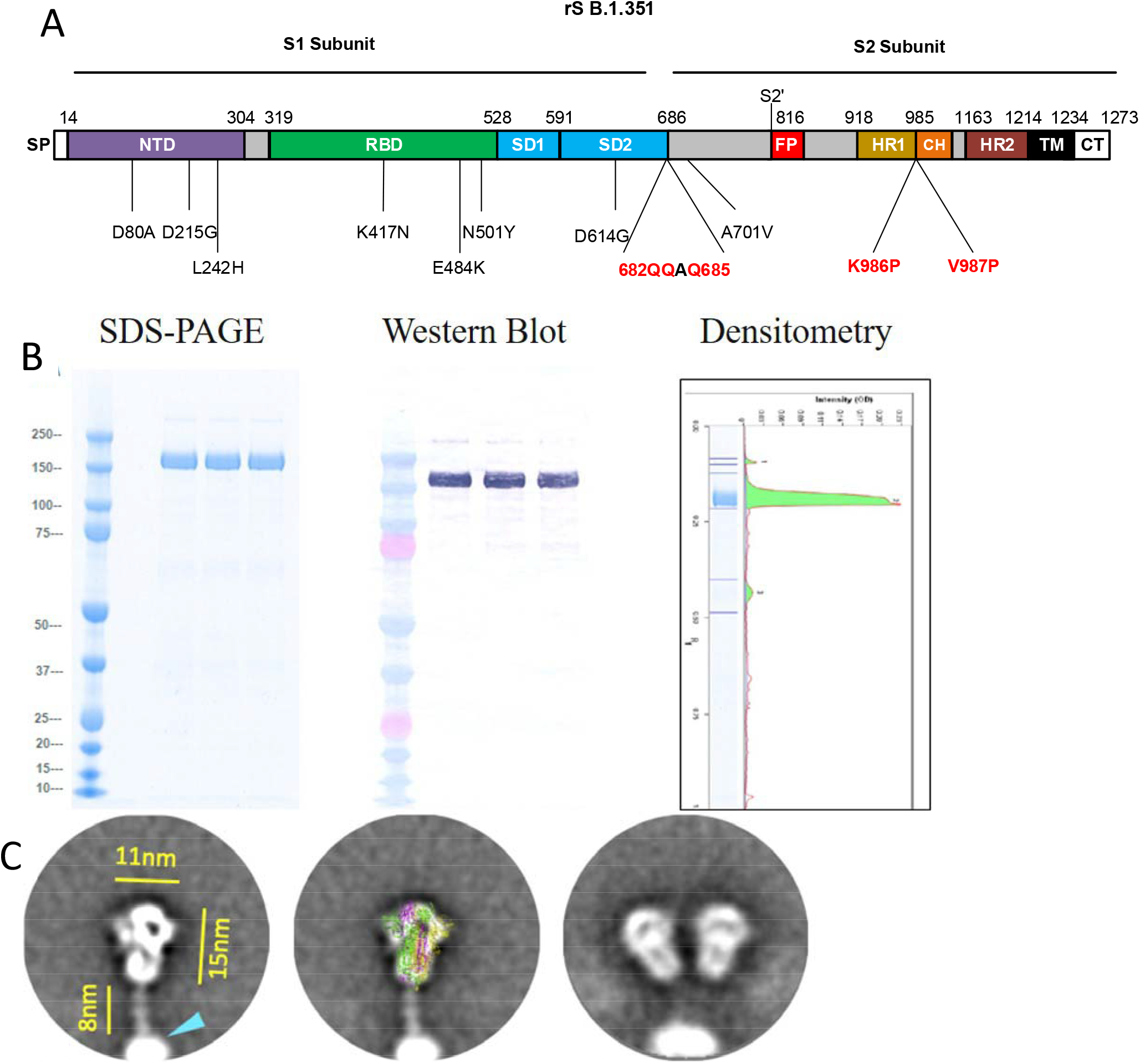
Characterization of SARS-CoV-2 Recombinant Spike Protein Construct Based on B.1.351 Variant. **A**. Linear diagram of the full-length SARS-CoV-2 spike (S) protein based on the protein sequence of the B.1.351 variant. Structural elements include the cleavable signal peptide (SP, white), N-terminal domain (NTD, purple), receptor binding domain (RBD, green), subdomains 1 and 2 (SD1 and SD2, blue), S2 cleavage site (S2⍰), fusion peptide (FP, red), heptad repeat 1 (HR1, yellow), central helix (CH, orange), heptad repeat 2 (HR2, brown), transmembrane domain (TM, black), and cytoplasmic tail (CT, white). Amino acid changes from the prototype rS protein sequence (rS-WU1) are shown in black text underneath the linear diagram. The native furin cleavage site was mutated (RRAR → QQAQ) to resist proteolytic cleavage and two proline mutations were also introduced to increase stability; these mutations are noted in red text underneath the linear diagram. **B**. Reduced SDS-PAGE gel with Coomassie blue staining of purified full-length rS-B.1.351 construct (left panel) showing the main protein product at the expected molecular weight of ∼170 kD. Western blot using a Spike-specific primary antibody confirming the identity of the main protein product (center panel). Scanning densitometry results are shown in the right panel. **C**. Negative stain transmission electron microscopy and 2D class averaging of rS-B.1.351. 2D images of rS-B.1.351 showed well-defined lightbulb-shaped particles with a length of 15nm and a width of 11nm. Trimers exhibited an 8nm flexible linker connected to PS-80 micelles as indicated with a cyan arrow in the left panel. Class average images showed a good fit of the rS-B.1.351 trimer with a cryo-EM solved structure of the prefusion SARS-CoV-2 trimeric spike protein ectodomain (PDB ID 6VXX) overlaid on the 2D image (middle panel). The right panel shows two rS-B.1.351 trimers anchored into a PS-80 micelle.

### Biophysical Properties, Structure, and Function of the rS-B.1.351 antigen

Purified SARS-CoV-2 recombinant spike (rS) construct rS-B.1.351, when reduced and subjected to SDS-PAGE, migrated with the expected molecular weight of approximately 170 kDa (Figure 1B). The thermal stability of rS-B.1.351 was compared to that of rS-WU1 by differential scanning calorimetry (DSC); the main peak of the rS-B.1.351 protein showed a 4°C increase in thermal transition temperature (T_max_) and 1.3-fold higher enthalpy of transition (ΔHCal) compared to the prototype rS-WU1 protein, indicating increased stability of the B.1.351 variant construct (Table 1). Negative stain transmission electron microscopy (TEM) imaging combined with two rounds of two-dimensional (2D) class averaging of 16,049 particles were used to confirm the ultrastructure of rS-B.1.351. High magnification (92,000x and 150,000x) TEM images revealed a lightbulb-shaped particle appearance with a 15nm length and an 11nm width, which was consistent with the prefusion form of the SARS-CoV-2 spike trimer (PDB ID 6VXX; Figure 1C). This is consistent with what we have previously observed for the prototype rS-WU1 spike protein (*14*).

**Table 1.** Thermostability and hACE2 binding of SARS-CoV-2 recombinant spike proteins.

| SARS-CoV-2 rS proteins | Differential Scanning Calorimetry (DSC) |  | hACE2 binding |  |  |
| --- | --- | --- | --- | --- | --- |
| | T<sub>max</sub> (°C) | $\Delta H_{cal}$ (kJ mol<sup>-1</sup>) | hACE2 Binding Kinetics by Bio-layer Interferometry | | hACE2 ELISA (EC<sub>50</sub>, ng/mL) |
|  |  |  | K<sub>a</sub> (1/Ms) | K<sub>dis</sub> (1/s) |  |
| rS-B.1.351 | 67.24 | 725.1 | $3.94 \times 10^4$ | $1.46 \times 10^{-7}$ | 8.0 |
| rS-WU1 | 63.21 | 546.0 | $1.08 \times 10^4$ | $1.56 \times 10^{-7}$ | 9.4 |

T<sub>max</sub>, melting temperature; K<sub>a</sub>, binding constant; K<sub>dis</sub>, dissociation constant; EC<sub>50</sub>, half-maximal binding.

To confirm the functional properties of the variant spike protein construct rS-B.1.351, binding of this rS to the hACE2 receptor was determined using bio-layer interferometry (BLI) as previously described (*14*). rS-B.1.351 was found to bind tightly and stably to hACE2, with an association constant (Ka) of 3.94 × 10^4^, representing a 3.6-fold greater association to hACE2 compared to the prototype protein rS-WU1 (Ka = 1.08 × 10^4^). Dissociation constants of these two proteins were essentially identical (1.46 × 10^−7^ and 1.56 × 10^−7^ for rS-B.1.351 and rS-WU1, respectively). We additionally assessed rS-B.1.351 binding to hACE2 with an ELISA as previously described (*14*). In this assay, rS-B.1.351 attained 50% saturation of hACE2 at a slightly lower concentration (EC_50_ = 8.0 ng/mL) than the prototype construct rS-WU1 (EC_50_ = 9.4 ng/mL), confirming that rS-B.1.351 variant spike exhibited a slightly higher affinity for hACE2 compared to that of rS-WU1 (Table 1).

### rS-B.1.351 vaccine immunogenicity in mice

We assessed the antibody- and cell-mediated immunogenicity of rS-B.1.351 and rS-WU1 (B.2 Spike) formulated with Matrix-M™ adjuvant. To assess antibody-mediated immunogenicity, groups of mice (n = 20) were immunized with either rS-WU1 or rS-B.1.351 as both prime and boost, with rS-WU1 as the prime and rS-B.1.351 as the boost, or with both vaccines combined in a bivalent formulation for the prime and boost vaccination. A placebo group received vaccine formulation buffer as a negative control. In monovalent immunization groups, 1 µg of rS and 5 µg of Matrix-M adjuvant was intramuscularly injected at Days 0 and 14. For bivalent immunization, 1 µg of each rS construct was administered at each immunization, for a total of 2 µg rS, with 5 µg of Matrix-M adjuvant. The study design is shown in Figure 2A. Mice immunized with either of the 4 vaccine regimens displayed elevated antibody titers against both the B.2 Spike and B.1.351 Spike by ELISA at day 21 post vaccination. Bivalent vaccination or heterologous vaccination produced significantly higher anti-S (WU1) IgG titers than monovalent vaccination with either rS-WU1 or rS-B.1.351, although group differences in IgG titers remained within 2-fold or less (Figure 2b). Regarding IgG titers against B.1.351 Spike, immunization with monovalent rS-B.1.351 or bivalent rS resulted in anti-B.1.351 spike IgG titers that were the highest among regimens tested, with no significant difference between these regimens. Immunization with rS-WU1 alone resulted in significantly lower titers compared to all other immunization regimens (Figure 2b). Animals in the placebo group exhibited undetectable anti-B.2 Spike and anti-B.1.351 spike IgG titers as expected.

**Figure 2.**
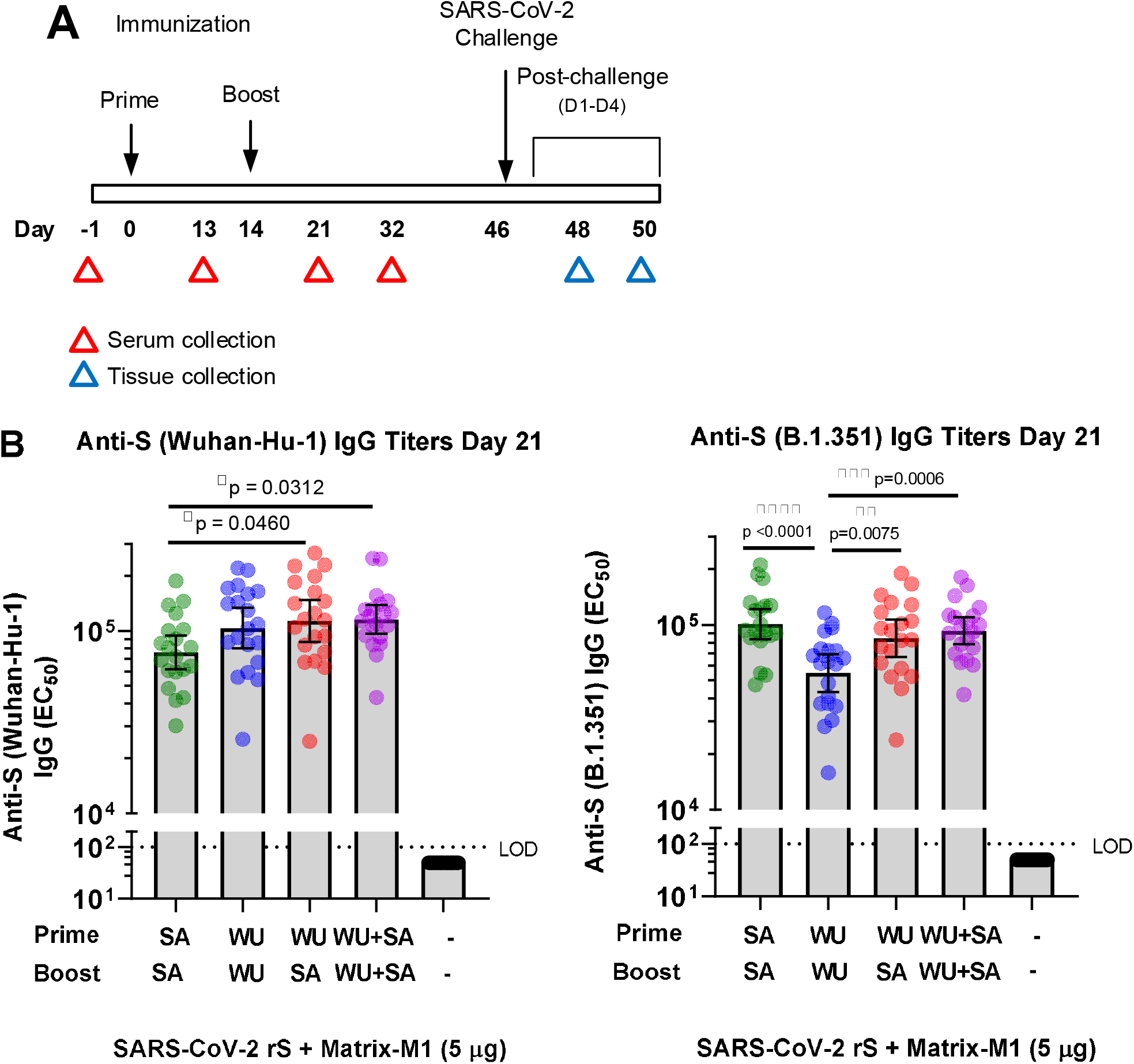
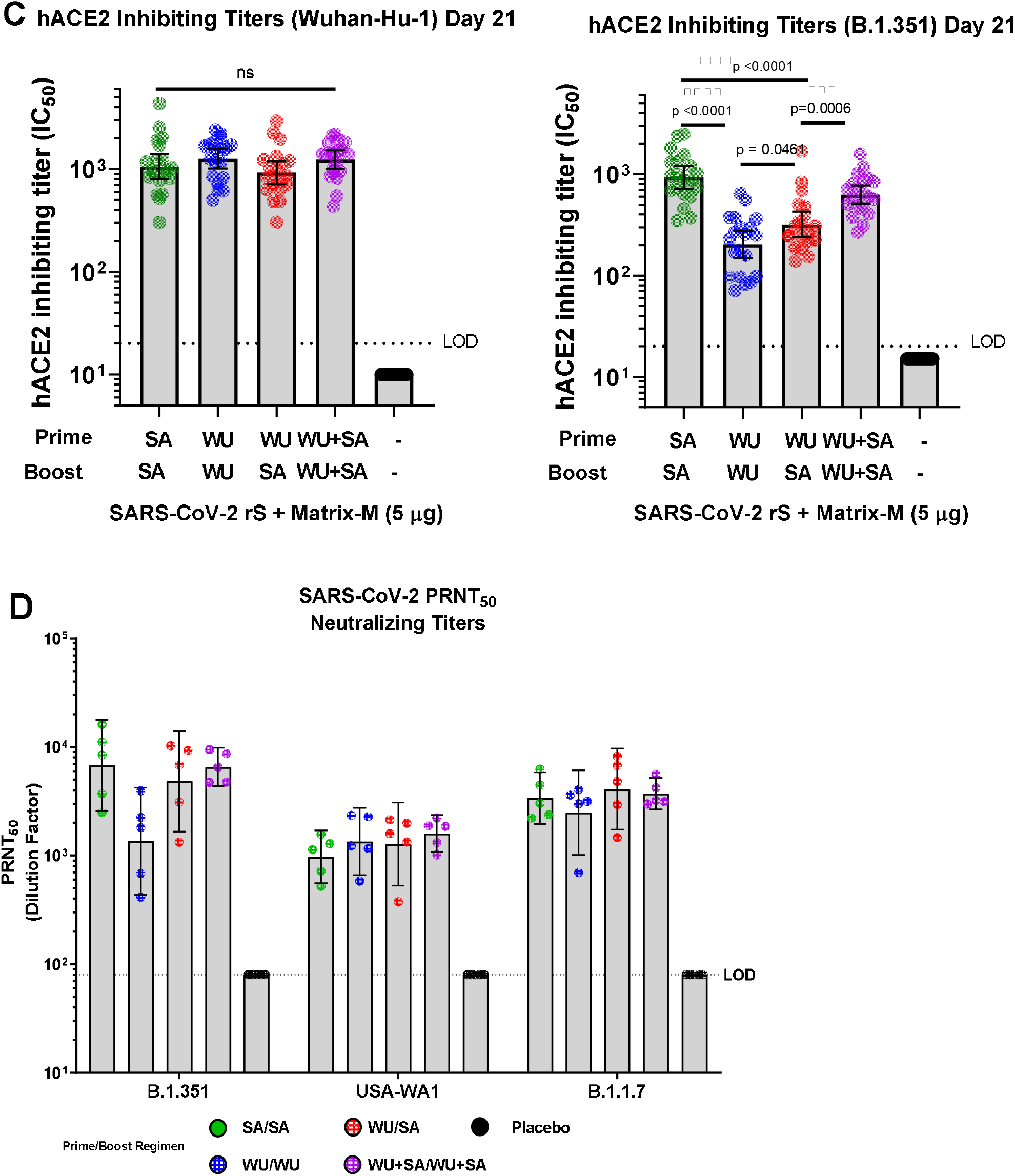
Antibody-mediated immunity induced upon immunization with SARS-CoV-2 rS based on Wuhan-Hu-1 or B.1.351 variant in mice. **A**. Groups of mice (n = 20/group) were immunized in a prime/boost regimen on study days 0 and 14 with various combinations of SARS-CoV-2 rS based on B.1.351 (SA) or Wuhan-Hu-1 (WU). Mice were either primed and boosted with rS-B.1.351, primed and boosted with rS-WU1, primed with rS-WU1 and boosted with rS-B.1.351, or primed and boosted with bivalent rS-WU1+rS-B.1.351. Antigen doses were 1 µg rS for each monovalent immunization, or 1 µg rS for each construct upon bivalent immunization (2 µg rS total). All antigen doses were administered with 5 µg Matrix-M1 adjuvant. A control group received formulation buffer (Placebo). Sera and tissues were collected at the timepoints listed in the diagram. **B**. Anti-SARS-CoV-2 S IgG serum titers were measured in sera collected on Day 21 using an ELISA to measure antibody titers against the Wuhan-Hu-1 spike protein (left panel) or B.1.351 spike protein (right panel). Bars indicate the geometric mean titer (GMT) and error bars represent 95% confidence interval (CI) for each group. Individual animal titers are indicated with colored symbols. **C**. ELISA was also used to determine the functional antibody titers in sera collected on Day 21 capable of disrupting binding between the SARS-CoV-2 receptor hACE2 and Wuhan-Hu-1 spike protein (left panel) or B.1.351 spike protein (right panel). Bars indicate the geometric mean titer (GMT) and error bars represent 95% confidence interval (CI) for each group. Individual animal titers are indicated with colored symbols. **D**. SARS-CoV-2 neutralizing antibody titers in sera collected on Day 32 from n = 5 animals/group were determined using a PRNT assay. Sera were evaluated for their ability to neutralize SARS-CoV-2 USA-WA1, B.1.351 variant, or B.1.1.7 variant. Bars indicate the geometric mean titer (GMT) and error bars represent 95% confidence interval (CI) for each group. Individual animal titers are indicated with symbols. Statistical significance was calculated by performing one-way ANOVA with Tukey’s post hoc test on log_10_-transformed data; significant differences among groups are indicated with asterisks and p-values in panels B and C.

The ability of serum from mice to inhibit Spike to hACE2 binding was also assessed (Figure 2c). All immunization regimens resulted in the production of antibodies that blocked hACE2 binding to B.2 spike with no significant difference between any groups at Day 21. Yet immunization with rS-WU1 alone resulted in significantly lower serum titers capable of disrupting binding between B.1.351 spike and hACE2; titers in the rS-B.1.351 alone immunization group were 4.6-fold higher than titers in the rS-WU1 alone immunization group (p < 0.0001) and titers in the group that received bivalent rS were 3.1-fold higher than titers in the rS-WU1 alone group (p < 0.0001). Monovalent rS-B.1.351 immunization or bivalent rS immunization also resulted in significantly higher titers than heterologous immunization (p < 0.0001 and p = 0.006, respectively).

We next assessed neutralizing antibody titers among the different vaccination regimens. Sera collected from vaccinated animals at day 32 post vaccination were assessed using SARS-CoV-2/WA1, SARS-CoV-2/B.1.1.7 and SARS-CoV-2/B.1.351 strains in a plaque reduction neutralizing titer assay (PRNT_50_, Figure 2d). Sera from the monovalent rS-WU1 group displayed similar neutralizing antibody titers to each of the 3 virus strains. Sera from mice immunized with monovalent rS-B.1.351 produced elevated neutralizing antibody titers to the B.1.351 and the B.1.1.7 strain compared to the B.2 strain. The heterologous vaccine group produced similar elevated neutralizing antibody titers to the B.1.351 and the B.1.1.7 strain compared to the B.2 strain, as did the bivalent rS-WU1/rS-B.1.351 vaccination regimen.

### rS-B.1.351 protection against SARS-CoV-2 in BALB/c mice

Mice vaccinated as described in Figure 2A were evaluated for their ability to produce protective immunity against challenge with either B.1.1.7 or B.1.351. While the SARS-CoV-2/Wuhan 1 (B.2) strain does not replicate in wild type mice, the B.1.1.7 and B.1.351 strains have a 501Y mutation in the Spike ORF allowing for Spike protein to bind to mouse ACE2 and enter cells (13)(*15*). At day 46 post vaccination, mice were intranasally inoculated with either 7 × 10^4^ PFU of B.1.1.7 (n = 10 mice per group) or 1 × 10^5^ PFU of B.1.351 (n = 10 mice per group). Mice were weighed daily throughout the post-challenge period, and at 2 and 4 days post infection (Study Days 48 and 50), 5 mice per group were euthanized by isoflurane inhalation. Lungs of each mouse were then assessed for viral load by plaque formation assay and viral RNA by RT-PCR. Placebo BALB/c mice infected with B.1.1.7 did not lose weight and there was also no observed weight loss in any vaccinated groups after being infected with this SARS-CoV-2 strain. For B.1.351 infected mice, 20% weight loss was observed in the placebo vaccination group by day 4 post infection with B.1.351 (Fig 3A). All mice vaccinated with either regimen were protected from weight loss after infection with B.1.351, demonstrating a clinical correlate of protection in this model.

**Figure 3.**
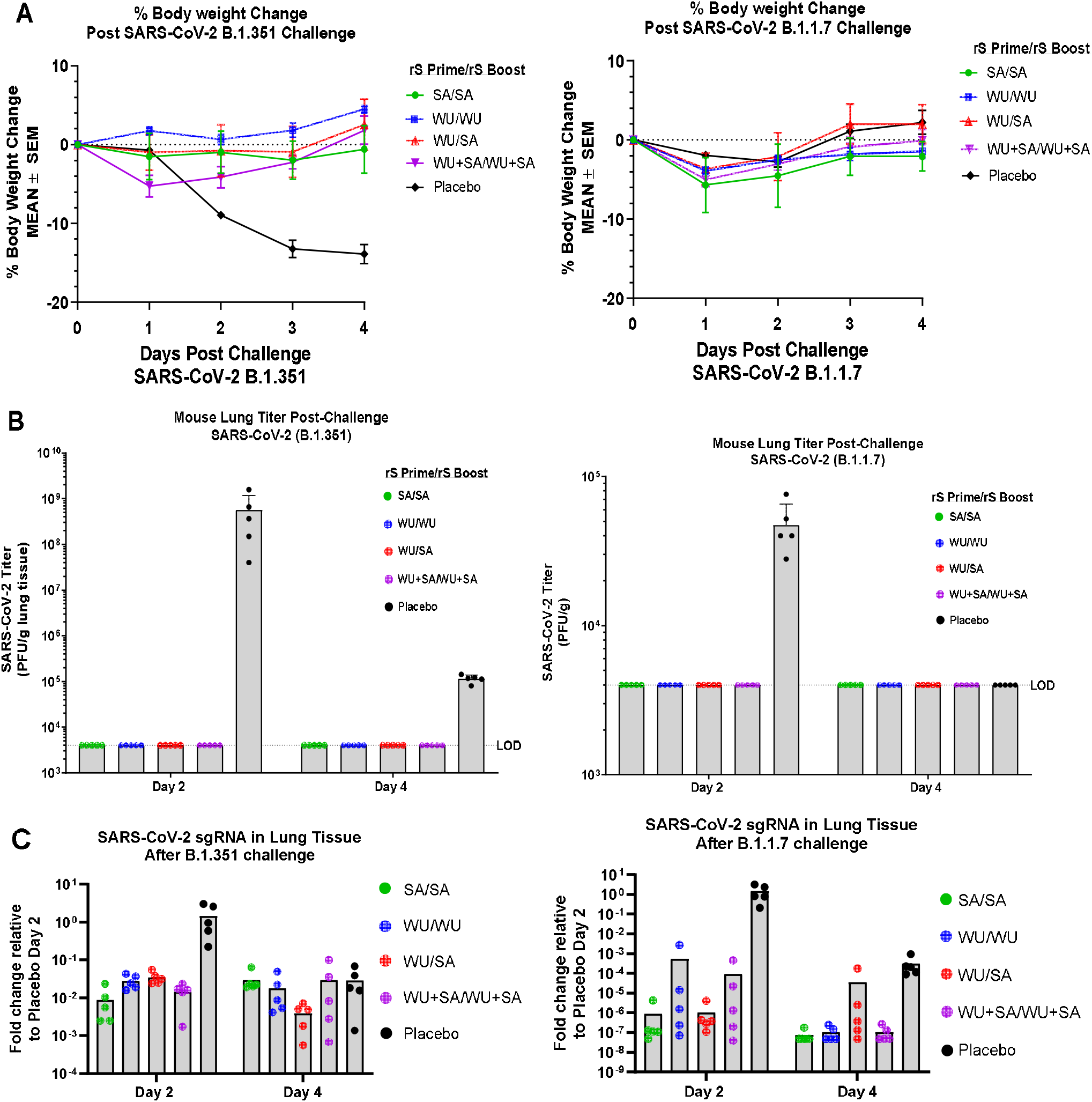
Protective efficacy of immunization with SARS-CoV-2 rS based on Wuhan-Hu-1 or B.1.351 against challenge with live SARS-CoV-2 B.1.351 or B.1.1.7 virus. Study design was described in Figure 2A. Immunized mice (n = 10/group) were challenged with live SARS-CoV-2 B.1.351 (left panels) or B.1.1.7 (right panels). For four days after challenge, mice were weighed daily and their percentage weight loss was calculated relative to their body weight on challenge day (**A**). Mean percentage body weight loss is shown with symbols and error bars represent standard error of the mean. Half of the mice were sacrificed at 2 days post-challenge and lung tissue was subjected to a plaque formation assay to determine lung viral titers, the remaining mice were sacrificed at 4 days post-challenge (**B**). Levels of SARS-CoV-2 subgenomic RNA were also determined in lung tissue and expressed as fold-change in RNA relative to the mean in the respective Placebo group on Day 2 post-challenge (**C**). Horizontal bars represent group mean fold-change from n = 5 mice at each timepoint and error bars represent standard deviation.

At day 2 post infection, B.1.1.7 infected mice in the placebo group exhibited 4 × 10^4^ pfu/g lung, which dropped to undetectable levels by day 4 post infection in the placebo vaccinated group. Upon immunization with any rS-WU1 or rS-B.1.351 regimen, there was no detectable live virus at day 2 or day 4 post infection, demonstrating a greater than 2-log reduction in viral load and protection from infection following vaccination (Fig 3B). At day 2 post infection, B.1.351 infected mice in the sham vaccinated group exhibited 8 × 10^8^ pfu/g lung, which dropped to 2 × 10^5^ pfu/g lung by day 4 post infection. Upon immunization with any rS regimen, there was no detectable live virus at day 2 or day 4 post infection in the B.1.351 infected mice. This demonstrates a dramatic reduction in virus titer, with > 5 log reduction in viral load by day 2 post infection from the sham vaccinated mice (Fig 3B). Lung RNA was also assayed for subgenomic (sgRNA) SARS-CoV-2 mRNA production after challenge. Relative to levels in the respective Placebo groups, we found >99% reduction in lung sgRNA levels in immunized mice at day 2 and day 4 after infection with both strains (Fig 3C).

These results confirm that rS-WU1 and rS-B.1.351 formulated with Matrix-M1 adjuvant and administered as monovalent, bivalent, or heterologous regimens confer protection against both strains of SARS-CoV-2, B.1.1.7 and B.1.351, in mice. Together with the reduction in weight loss, high neutralizing antibody titers, and elimination of viral replication in the lungs of mice, we demonstrate a highly protective vaccine response by the variant Spike targeted vaccine.

### Cell-mediated immunogenicity of rS-B.1.351 in mice

Groups of BALB/c mice (n = 8/group) were immunized with the same rS-WU1 or rS-B.1.351 regimens mentioned above, but at a 21-day interval. A negative control group (n = 5) was injected with vaccine formulation buffer. Spleens were harvested on day 28, 7 days after the boost immunization. Splenocytes were collected and subjected to ELISpot and intracellular cytokine staining (ICS) to examine cytokine secretion upon stimulation with rS-WU1 or rS-B.1.351. ELISpot assay showed greater numbers of IFN-γ producing cells compared to the number of IL-5 producing cells upon all vaccination regimens, signifying a Th1-skewed response (Fig 4B-D). Upon stimulation with either rS, strong Th1 responses were observed by ICS as measured by the presence of CD44^hi^CD62^low^CD4^+^ effector memory T cells expressing IFN-γ, IL-2, or TNF-α, and multifunctional CD4^+^ T cells expressing all three cytokines (Figure 4E, Figure S1). Th2 CD4^+^ T cells that expressed IL-4 but were negative for IL-2 and TNF-α were also present, but at a lower proportion than that observed for Th1 cytokines (Figure S1). No significant differences in cytokine-positive cell number were observed among vaccination groups for any cytokine tested upon stimulation with either rS-WU1 or rS-B.1.351.

**Figure 4.**
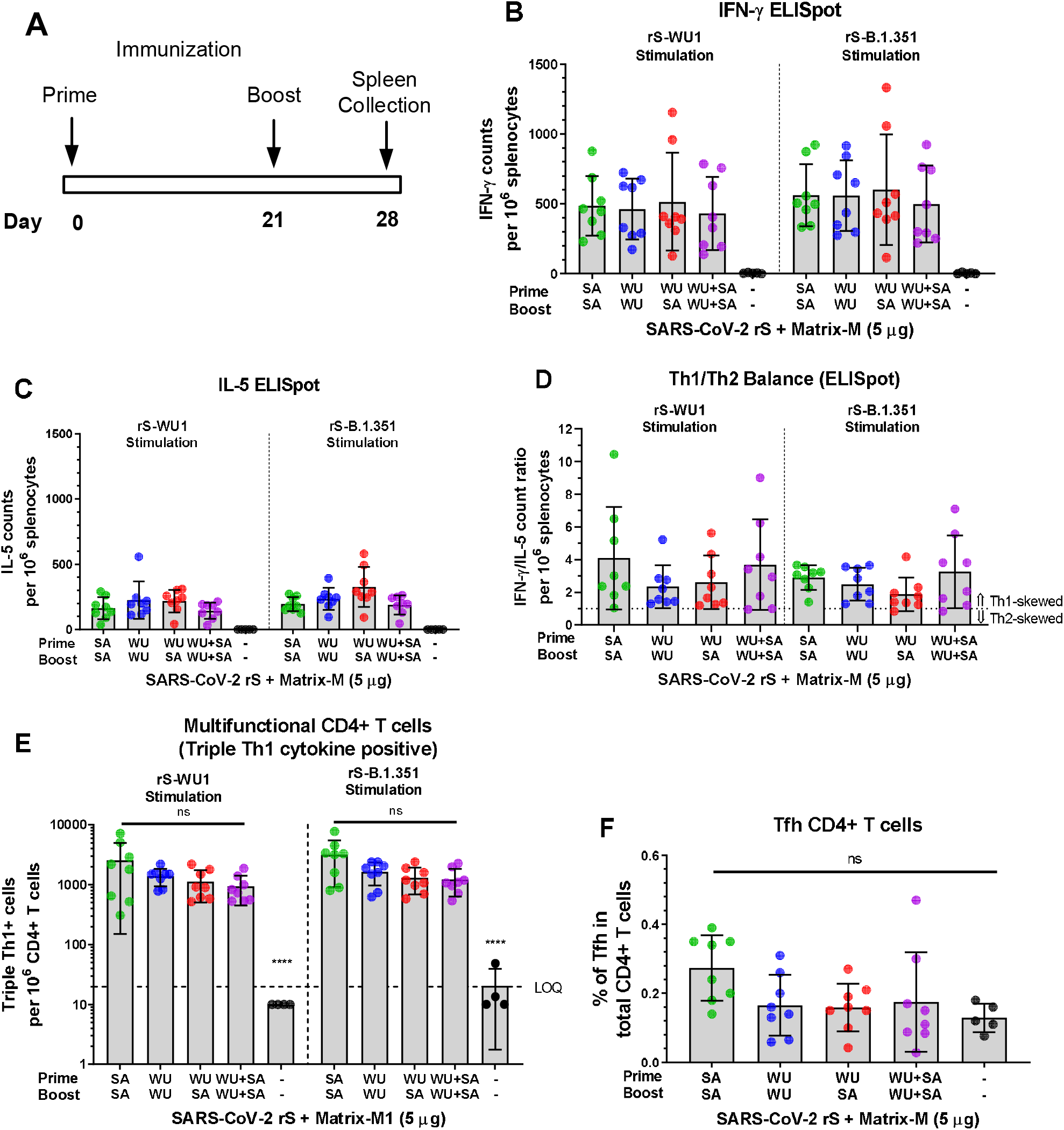
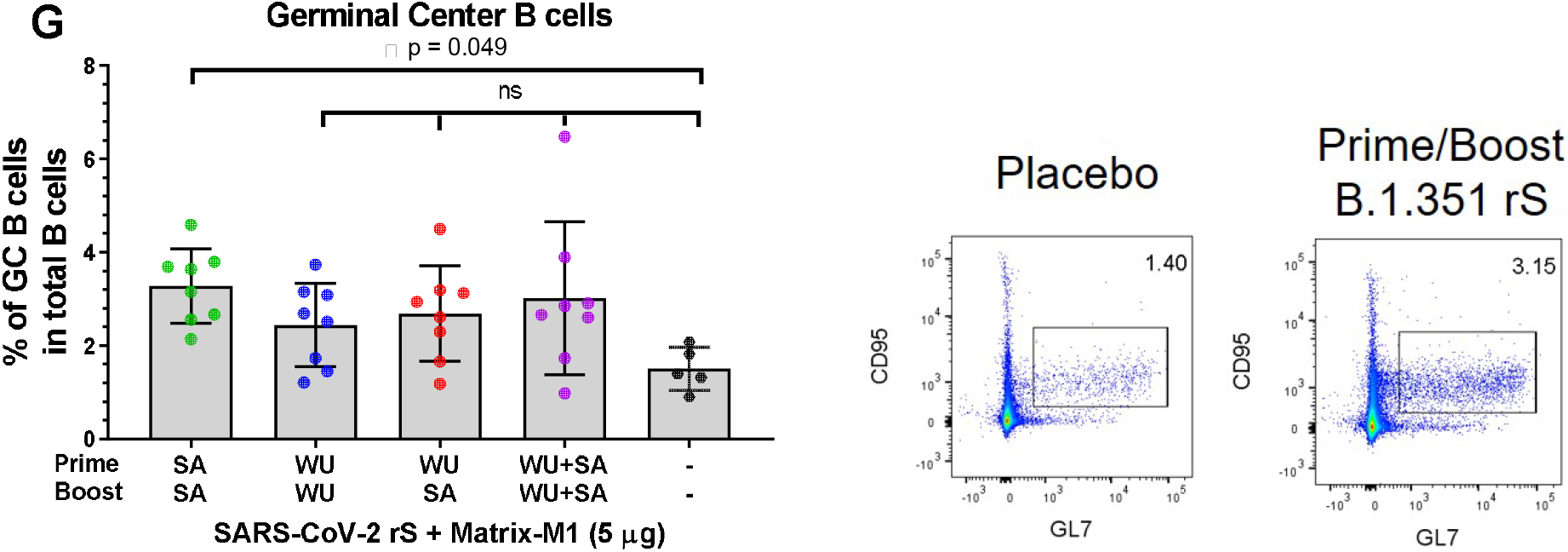
Cell-mediated immunity upon immunization with rS-WU1 or rS-B.1.351 regimens in mice. **A.** Groups of mice (n = 8/group) were immunized in a prime/boost regimen on Days 0 and 21 with various combinations of SARS-CoV-2 rS based on B.1.351 (SA) or Wuhan-Hu-1 (WU). Mice were either primed and boosted with rS-B.1.351, primed and boosted with rS-WU1, primed with rS-WU1 and boosted with rS-B.1.351, or primed and boosted with bivalent rS-WU1+rS-B.1.351. Antigen doses were 1 µg rS for each monovalent immunization, or 1 µg rS for each construct upon bivalent immunization (2 µg rS total). All immunizations were administered with 5 µg Matrix-M1 adjuvant. A control group received formulation buffer (Placebo, n = 5). Spleens were harvested on Day 28 for cell collection. Splenocytes were stimulated with rS-WU1 or rS-B.1.351, then subjected to ELISpot assay to determine IFN-γ-positive cells as a representative Th1 cytokine **(B)** and IL-5-positive cells as a representative Th2 cytokine **(C)**. Data from panels B and C were used to calculate the Th1/Th2 balance of responses to immunization **(D). E**. The numbers of multifunctional CD4+ T cells that stained positively for three Th1 cytokines (IFN-γ, IL-2, and TNF-α) using intracellular cytokine staining were quantified and expressed as the number of triple cytokine positive cells per 10^6^ CD44^hi^CD62^Low^ effector memory CD4+ T cells. **F**. T follicular helper cells were quantified by determining the percentage of PD-1+CXCR5+ cells among all CD4+ T cells. **G**. Germinal center formation was evaluated by determining the percentage of GL7+CD95+ cells among CD19+ B cells using flow cytometry. Gray bars represent means and error bars represent standard deviation. Individual animal data are shown with colored symbols. An example of the gating strategy is shown in the right panel. Differences among experimental groups were evaluated by one-way ANOVA with Tukey’s post-hoc test (data in panel B were log_10_-transformed before analysis). P values < 0.05 were considered statistically significant; **** = p < 0.0001.

T follicular helper cells (CXCR5^+^PD-1^+^CD4^+^) represent a group of CD4^+^ T cells in the lymphoid organs that aid germinal center formation and B cell development. We observed a trend in expansion of T follicular cells in the vaccinated groups, though no statistically significant elevation was observed compared to placebo animals (Figure 4F). Similarly, germinal center formation was evaluated by determining the percentage of GL7^+^CD95^+^CD19^+^ germinal center B cells among total B cells using flow cytometry, and though a tendency toward higher percentage of germinal center B cells was observed in vaccinated groups compared to the placebo group, only animals immunized with the monovalent rS-B.1.351 regimen showed a significantly higher proportion (p = 0.049 compared to placebo; Figure 4G).

### Anamnestic response induced by boosting with rS-B.1.351 one year after primary immunization with rS-WU1 in baboons

A small cohort of baboons (n = 9 total) were subjected to a primary immunization series with rS-WU1 (either 1 µg, 5 µg, or 25 µg rS with 50 µg Matrix-M adjuvant, or unadjuvanted 25 µg rS). Approximately one year later, all animals were boosted with one or two doses of 3 µg rS-B.1.351 with 50 µg Matrix-M adjuvant to examine the resulting immune responses (Figure 5A). Seven days after the first rS-B.1.351 boost, animals that had originally received adjuvanted rS-WU1 exhibited a strong anamnestic response as exhibited by levels of anti-S (WU1) IgG titers higher than that originally observed at peak immune response during the primary immunization series (Figure 5B). This response did not seem to be further bolstered by a second booster dose of rS-B.1.351, though the small sample sizes utilized in this study prohibit a meaningful quantitative analysis. Animals that received unadjuvanted rS-WU1 during the primary immunization series exhibited a weaker response to boosting with rS-B.1.351, though still exhibited elevated anti-S (WU1) IgG response. The rS-B.1.351 boost elicited comparable antibody titers against rS-WU1 and rS-B.1.351, with animals that originally received unadjuvanted rS-WU1 exhibiting a weaker response (Figure 5C).

**Figure 5.**
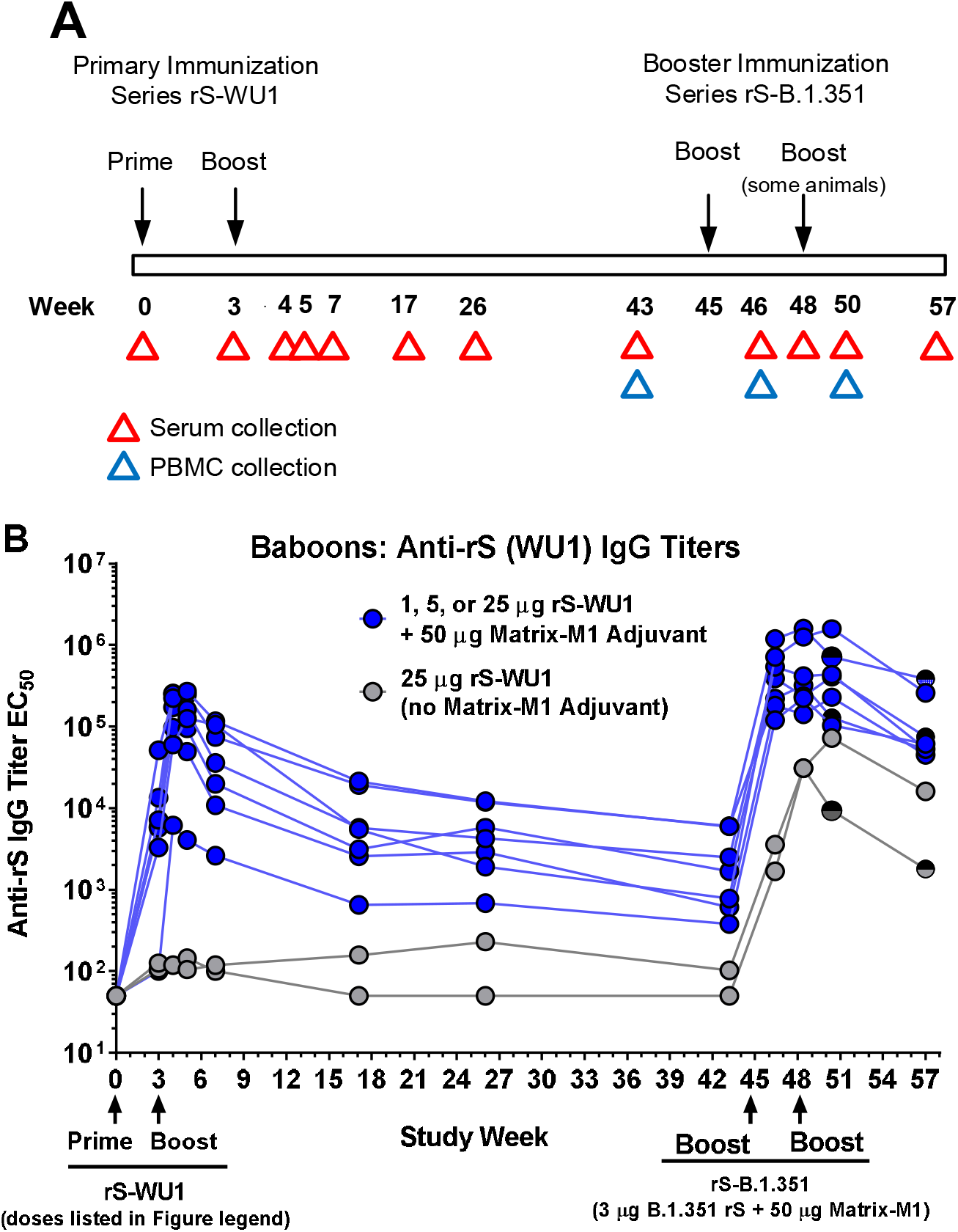
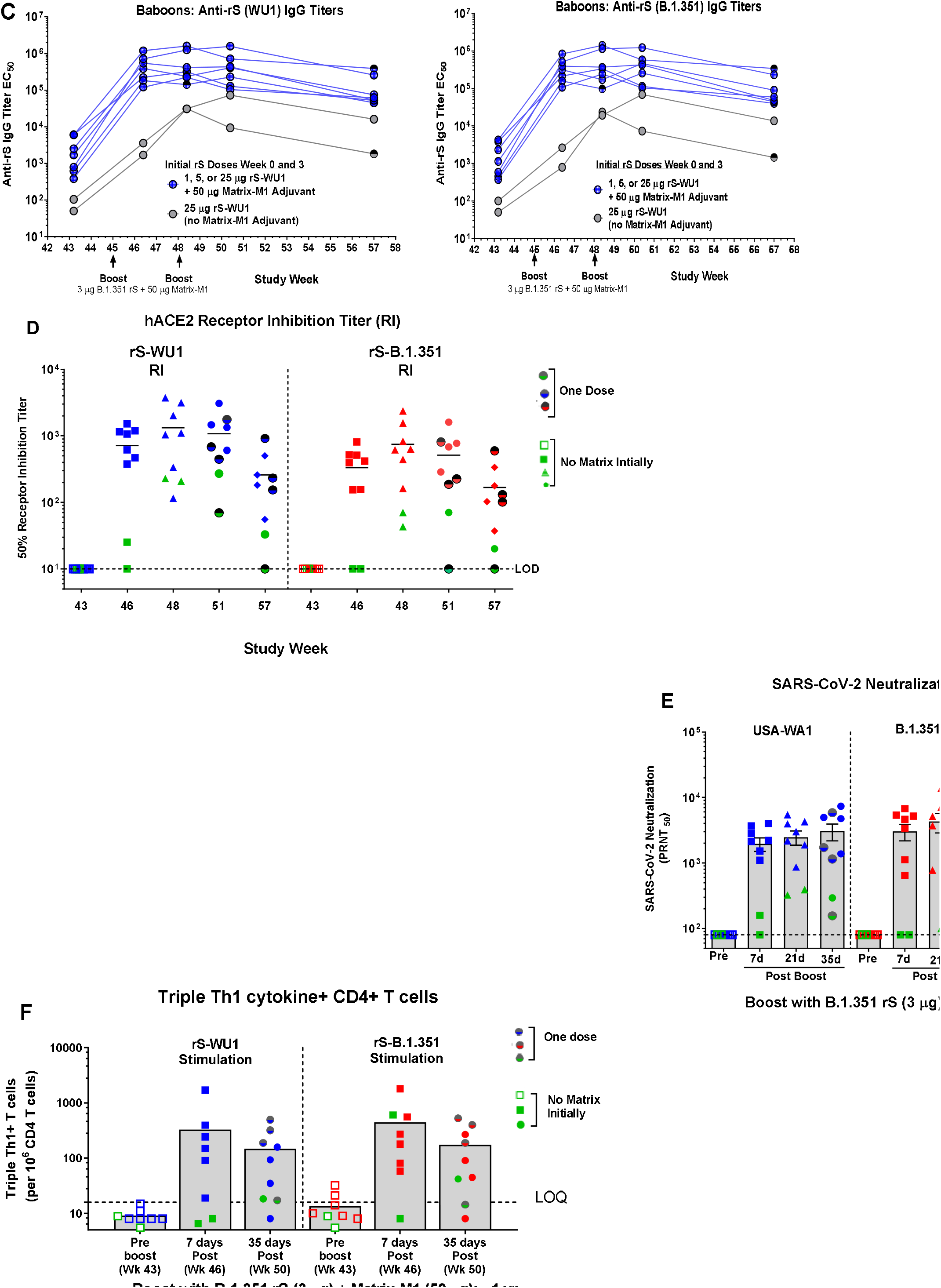
Immunogenicity of one or two rS-B.1.351 booster doses approximately one year after immunization with rS-WU1 in baboons. **A**. A small cohort of baboons (n = 2-3/group) were originally immunized with 1 µg, 5 µg, or 25 µg rS-WU1 with 50 µg Matrix-M1 adjuvant or unadjuvanted 25 µg rS-WU1 on Day 0 and 21 (Week 0 and 3, respectively). Approximately 1 year later, all animals were boosted with one or two doses of 3 µg rS-B.1.351 with 50 µg Matrix-M1 adjuvant on Day 318 and 339 (Week 45 and 48, respectively). **B**. Anti-S (WU1) IgG titers were measured throughout the course of the study. Individual animals’ titers are shown over time, different colored symbols and lines represent different dose groups for the initial rS-WU1 immunization series. Sera collected pre-rS-B.1.351 boost (Day 303) as well as 7, 21, 35, and 81 days after the boost were analyzed to determine anti-rS-WU1 and rS-B.1.351 IgG titers by ELISA (**B**, horizontal lines represent means), antibody titers capable of disrupting the interaction between rS-WU1 or rS-B.1.351 and the hACE2 receptor by ELISA (**D**, horizontal lines represent means), and antibody titers capable of neutralizing SARS-CoV-2 strains USA-WA1, B.1.351, and B.1.1.7 with a PRNT assay (**E**, gray bars represent geometric means and error bars represent 95% confidence intervals). The presence of multifunctional CD4+ T cells positive for three Th1 cytokines (IFN-γ, IL-2, and TNF-α) was evaluated with intracellular cytokine staining after stimulation with rS-WU1 or rS-B.1.351 **(F)**. Gray bars represent means and colored symbols represent individual animal data.

Serum antibody titers capable of disrupting the interaction between B.2 rS or B.1.351 rS and hACE2 were also evaluated at before boost, and 7, 21, 35, and 89 days after the boost with 1 or 2 doses of rS-B.1.351. Similarly, to what was observed for anti-S IgG titers, animals that had received adjuvanted vaccine during the primary immunization series exhibited a strong hACE2-inhibiting antibody response 7 days after the rS-B.1.351 boost, despite having undetectable titers before the boost. Titers were slightly higher for rS-WU1-hACE2 blocking antibodies compared to levels of rS-B.1.351-hACE2 blocking antibodies, though the small sample size prohibits a meaningful quantitative analysis. Animals that had received unadjuvanted vaccine during the primary immunization series exhibited lower hACE2 blocking titers after the rS-B.1.351 boost (Figure 5D).

Neutralizing antibody titers were analyzed by Plaque Reduction Neutralization Titer (PRNT) assay by testing sera for the ability to neutralize WA1, B.1.351 and B.1.1.7. Sera collected just before the rS-B.1.351 boost had undetectable neutralizing antibody levels against all these viruses. By 7 days post vaccination, high titer antibody that neutralized all 3 strains was detected and this antibody response stayed high through 35 days post vaccination. Animals immunized with unadjuvanted rS-WU1 in the primary series displayed significantly lower antibody levels with a much broader range of neutralization titers (Figure 5E). Together, these data demonstrate a robust durable antibody response even 1 year after the primary vaccination series.

Multifunctional CD4^+^ T cells expressing three Th1 cytokines were also expanded 7 days after the first rS-B.1.351 booster dose in baboons, and these responses were maintained at 35 days after the first booster dose (Figure 5F and Figure S2).

### Neutralization of SARS-CoV-2 variants by sera from NVX-CoV2373 vaccinated adults

NVX-CoV2373 is currently in clinical trials globally, including in locations where B.1.1.7 and B.1.351 are prevalent. We assessed the capacity of sera from individuals in these trials to neutralize USA-WA1, B.1.1.7 and B.1.351 by performing a PRNT_50_ (Figure 6). Thirty randomly selected human serum samples from Phase 2 clinical trial participants after their second dose of the vaccine were assayed. When comparing WA1 vs B.1.1.7, there was no change in neutralizing activity across the majority of the serum samples; only 1 sample had a statistically significant change in neutralizing antibody titers against B.1.1.7. The WA1 vs B.1.351 neutralization titers showed increased range of neutralization titers with five out of 30 samples showing reduced neutralization 1 standard deviation away from the mean in the PRNT_50_ assay. These data demonstrate a reduced neutralization of B.1.351 in a small percentage of NVX-CoV2373 vaccinees compared to B.1.1.7. These data support the development and production of a B.1.351 targeted vaccine, which we demonstrated was efficient at protecting mice against B.1.351 challenge and can induce a strong anamnestic response in NHPs originally immunized with prototype vaccine.

**Figure 6.**
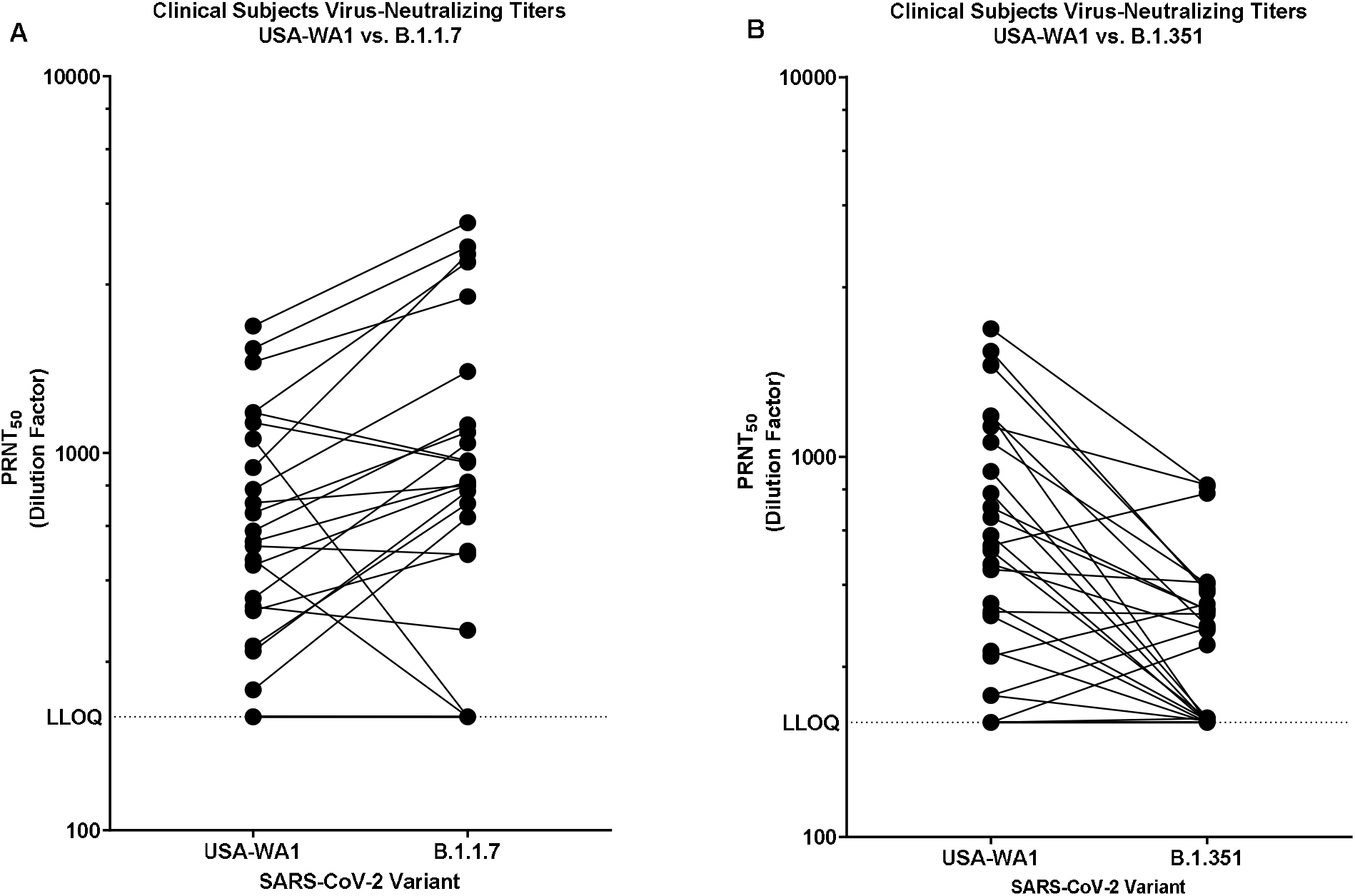
SARS-CoV-2 variant neutralizing titers from human subjects immunized with NVX-CoV2373. Serum samples from clinical study participants (n = 30, 2019nCoV-101-P2) were tested in a PRNT assay to determine the presence of neutralizing antibodies against USA-WA1 compared to B.1.1.7 **(A)** and B.1.351 variants **(B)**. Individual subjects’ titers are shown with black circles, lines connect individuals’ titers against USA-WA1 to their titer against the respective variant.

## Discussion

As SARS-CoV-2 has spread, it has created selective pressure for viruses with increased fitness across multiple parameters including replication rate, receptor binding, and antibody evasion (*16, 17*). This has produced several variant mutants that are gaining prevalence around the world. One of these variants, B.1.351, was first identified in South Africa. This variant has gained the ability to evade antibodies produced during both convalescence and vaccination and significantly evade currently authorized monoclonal antibody therapies. Based on the potential need for a spike variant-directed vaccine, we have produced a recombinant SARS-CoV-2 spike protein antigen based on the B.1.351 lineage (rS-B.1.351). We have shown that a full-length, stabilized prefusion SARS-CoV-2 spike glycoprotein vaccine using the B.1.351 Spike variant adjuvanted by Matrix-M can induce high levels of functional immunity and protects mice against both B.1.1.7 and B.1.351 SARS-CoV-2 strains. Immunizing mice or non-human primates with rS-B.1.351 induced anti-S antibodies, hACE2-receptor inhibiting antibodies, and SARS-CoV-2 neutralizing antibodies. In addition, the rS-B.1.351 vaccine induced Ag-specific CD4^+^ T cell responses, induced germinal center formation and provided protection against B.1.351 and B.1.1.7 challenge.

In mice, the antibodies produced after vaccination with the B.1.351 variant-directed vaccine were able to inhibit binding between hACE2 and variant spike or ancestral spike to the same degree, indicating that this variant-directed vaccine could efficiently protect “backward” against ancestral SARS-CoV-2 strains. In contrast, the prototype vaccine was less efficient at protecting “forward” against the B.1.351 variant. This finding corroborates previous findings that emerging strains of the coronavirus 229E can evade antibody immunity (16(*18*)), enforcing the need for variant-directed SARS-CoV-2 vaccines during primary immunization or as booster vaccines. The results in NHPs presented in this work suggest that one dose of a variant-directed vaccine may be sufficient for boosting regimens after previous immunization with a COVID-19 vaccine based on ancestral spike strains; two variant vaccine boosters did not seem to confer any added benefit over a single booster. Production of high neutralizing antibody titers after initial convalescence or after single vaccination points to the affinity maturation that occurs after first exposure to a new antigen. This immunological training may be essential for durability.

As variant viruses continue to emerge across the world, they are being selected for in the background of increasing seroprevalence. Vaccines must be able to produce high antibody titers across a wide range of age groups and comorbidities and the vaccine platform must be able to rapidly respond to the variants at the earliest signs of their emergence. The baculovirus-Sf9 cell core technology, used to produce rS protein, is highly amenable to a rapid response to emergent SARS-CoV-2 variants of concern; producing a variant-directed recombinant spike antigen requires few to no changes in the purification and manufacturing processes developed for producing prototype spike vaccines. After a prototype vaccine is shown to meet critical immunogenicity, efficacy, and safety benchmarks, variant-directed vaccines can also reach the clinic on an accelerated timeline from a regulatory standpoint. Analysis of human vaccine sera from our trials demonstrated a robust antibody response and minimal loss of neutralization. We observed that B.1.351 virus does not significantly reduce neutralization compared to B.1.1.7 and WA1, even though there is evidence of breakthrough infections in the WA1 trial participants in South Africa. All breakthrough infections were B.1.351 in this Phase 2b trial (11). This is suggestive of the broader findings that while most respond strongly to vaccination, there is a range of human antibody responses to any vaccine. A booster vaccination containing a single or multiple variant rS vaccines could both increase antibody levels as well as broaden coverage to variants as shown in this work.

## Methods

### Cells and Virus

Virus and cells were processed as described previously (*19*). Briefly, Vero E6 cells (ATCC# CRL 1586) were cultured in DMEM (Quality Biological), supplemented with 10% (v/v) fetal bovine serum (Gibco), 1% (v/v) penicillin/streptomycin (Gemini Bio-products) and 1% (v/v) L-glutamine (2 mM final concentration, Gibco) (Vero media). Cells were maintained at 37°C and 5% CO_2_. SARS-CoV-2/WA1 were provided by the CDC (BEI #NR-52281). SARS-CoV-2/B.1.17 and SARS-CoV-2/B.1.351 were generously provided by Dr. Andy Pekosz at The Johns Hopkins University. Stocks for both viruses were prepared by infection of Vero/TMPRSS2 cells for two days when CPE was starting to be visible(*20*). Media were collected and clarified by centrifugation prior to being aliquoted for storage at -80°C. Titer of stock was determined by plaque assay using Vero E6 cells as described previously(*21*). All work with infectious virus was performed in a Biosafety Level 3 laboratory and approved by our Institutional Biosafety Committee.

#### SARS-CoV-2 protein expression

SARS-CoV-2 constructs were synthetically produced from the full-length S glycoprotein gene sequence (GenBank MN908947 nucleotides 21563-25384). The full-length *S-*genes were codon optimized for expression in *Spodoptera frugiperda* (Sf9) cells and synthetically produced by GenScript® service (GenScript USA, Piscataway, NJ, USA). The QuikChange® Lightning site-directed mutagenesis kit (Agilent Technologies, Inc.) was used to produce two spike protein variants: the furin cleavage site (682-RRAR-685) was mutated to 682-QQAQ-685 to be protease resistant and two proline substitutions at positions K986P and V987P (2P) were introduced to produce the double mutant, BV2373. To generate the recombinant Spike construct based on the B.1.351 variant, the following point mutations were also introduced: D60A, D215G, L242H, K417N, E484K, N501Y, D614G, and A701V. Full-length *S*-genes were cloned between the BamHI – HindIII sites in the pFastBac baculovirus transfer vector (Invitrogen, Carlsbad, CA) under transcriptional control of the *Autographa californica* polyhedron promoter. Recombinant baculovirus constructs were plaque purified and master seed stocks prepared and used to produce the working virus stocks. The baculovirus master and working stock titers were determined using rapid titer kit (Clontech, Mountain View, CA). Recombinant baculovirus stocks were prepared by infecting Sf9 cells with a multiplicity of infection (MOI) of ≤0.01 plaque forming units (pfu) per cell.

#### Expression and purification

SARS-CoV-2 rS proteins were produced in Sf9 cells as previously described (*14*). Briefly, cells were expanded in serum-free medium and infected with recombinant baculovirus. Cells were cultured at 27 ± 2°C and harvested at 68-72 hours post-infection by centrifugation (4000 × *g* for 15 min). Cell pellets were suspended in 25 mM Tris HCl (pH 8.0), 50 mM NaCl and 0.5-1.0% (v/v) TERGITOL NP-9 with leupeptin. rS proteins were extracted from the plasma membranes with Tris buffer containing NP-9 detergent, clarified by centrifugation at 10,000 × *g* for 30 min. rS proteins were purified by TMAE anion exchange and lentil lectin affinity chromatography. Hollow fiber tangential flow filtration was used to formulate the purified spike protein at 100-150 μg mL^-1^ in 25 mM sodium phosphate (pH 7.2), 300 mM NaCl, 0.02% (v/v) polysorbate 80 (PS 80). Purified rS proteins were evaluated by 4-12% gradient SDS-PAGE stained with Gel-Code Blue reagent (Pierce, Rockford, IL) and purity was determined by scanning densitometry using the OneDscan system (BD Biosciences, Rockville, MD).

#### Differential scanning calorimetry (DSC)

Samples (BV2426 Lot 01Feb21 and BV2373 Lot 15Dec20; rS-B.1.351 and rS-WU1, respectively) and corresponding buffers were heated from 4°C to 120°C at 1°C per minute and the differential heat capacity change was measured in a NanoDSC (TA Instruments, New Castle, DE). A separate buffer scan was performed to obtain a baseline, which was subtracted from the sample scan to produce a baseline-corrected profile. The temperature where the peak apex is located is the transition temperature (T_max_) and the area under the peak provides the enthalpy of transition (ΔHcal).

#### Transmission electron microscopy (TEM) and 2D class averaging

Electron microscopy was performed by NanoImaging Services (San Diego, CA) with a Thermo Fisher Scientific Glacios Cryo Transmission Electron Microscope, operated at 200kV equipped with a Falcon 3 direct electron detector. SARS-CoV-2 rS proteins were diluted to 16 µg mL^-1^ in formulation buffer. The samples (3 µL) were applied to continuous carbon grids and stained with uranyl format. Images of each grid were acquired at multiple scales to assess the overall distribution of the sample. High-magnification images were acquired at nominal magnifications of 150,000x (0.095 nm/pixel) and 92,000x (0.159 nm/pixel). The images were acquired at a nominal defocus of - 2.0µm to -1.5µm and electron doses of ∼25 e^-^/Å^2^.

For class averaging, particles were identified from 92,000x high magnification images, followed by alignment and classification as previously described(*14*).

#### Kinetics of SARS-CoV-2 S binding to hACE2 receptor by BLI

rS protein receptor binding kinetics was determined by bio-layer interferometry (BLI) using an Octet QK384 system (Pall Forté Bio, Fremont, CA). His-tagged human ACE2 (2 μg mL^-1^) was immobilized on nickel-charged Ni-NTA biosensor tips. After baseline, SARS-CoV-2 rS protein solutions were 2-fold serially diluted in kinetics buffer over a range of 300 nM to 4.7 nM, allowed to associate for 600 sec, followed by dissociation for an additional 600-900 sec. Data was analyzed with Octet software HT 10.0 by 1:1 global curve fit.

#### Animal ethics statement

The mouse immunizations were performed by Noble Life Sciences (Sykeville, MD) before transfer to UMSOM BSL3 facility. Noble Life Sciences and University of Maryland School of Medicine are accredited by the Association for Assessment and Accreditation of Laboratory Animal Care (AAALACC International). All animal procedures were in accordance with NRC Guide for the Care and Use of Laboratory Animals, the Animal Welfare Act, and the CDC/NIH Biosafety in Microbiological and Biomedical Laboratories. Mouse challenge studies were approved by The University of Maryland School of Medicine IACUC. The olive baboon (*Papio cynocephalus anubis*) study was performed at the University of Oklahoma Health Science Center (OUHSC). OUHSC is accredited by AAALACC International. Baboons were maintained and treated according to the Institutional Biosafety Committee guidelines. Baboon experiments were approved by the Institutional Animal Care and Use Committee (IACUC) and the Institutional Biosafety Committee of OUHSC. Studies were conducted in accordance with the National Institutes of Health Guide for Care and Use of Laboratory Animals (NIH publication 8023, Revised 1978).

#### Mouse study designs

Female BALB/c mice (7-9 weeks old, 17-22 grams, N = 20 per group) were immunized by intramuscular (IM) injection with two doses spaced 14 days apart (study day 0 and 14) of rS-WU1, rS-B.1.351 with 5 μg saponin-based Matrix-M™ adjuvant (Novavax, AB, Uppsala, SE) either alone, in combination, or as a heterologous prime/boost. A placebo group was injected with vaccine formulation buffer as a negative control. Serum was collected for analysis on study days -1, 14, 21, and 32. Vaccinated and control animals were intranasally challenged with SARS-CoV-2 on study day 46.

To assess the cellular response mediated by Matrix-M adjuvant, groups of female BALB/c mice (N = 8 per group) were immunized IM with the same regimens described above, with injections spaced 21 days apart. Spleens were collected 7 days after the second immunization (study day 28). A non-vaccinated group (N = 5) served as a control.

#### Baboon study design

Nine adult baboons (10-16 years of age at study initiation) were randomized into 4 groups of 2-3/group and immunized by IM injection with rS-WU1 at 1, 5, or 25 μg rS with 50 μg Matrix-M adjuvant. A separate group was immunized with 25 μg rS without adjuvant. Animals were vaccinated with 2 doses spaced 21 days apart in this primary immunization series. Immunogenicity results after the primary immunization series were previously described (*14*). Approximately one year later (45 weeks), all animals were boosted with one or two 3 μg doses of rS-B.1.351 with 50 μg Matrix-M adjuvant. Sera and PBMCs were collected before and after the boost to measure antibody- and cell-mediated immune responses.

#### SARS-CoV-2 challenge in mice

Mice were anaesthetized by intraperitoneal injection 50 μL of a mix of xylazine (0.38 mg/mouse) and ketamine (1.3 mg/mouse) diluted in phosphate buffered saline (PBS). Mice were intranasally inoculated with either 7 × 10^4^ pfu of B.1.1.7 or 1 × 10^5^ pfu of B.1.351 strains of SARS-CoV-2 in 50 μL. Challenged mice were weighed on day of infection and daily for 4 days post infection. At days 2 and 4 post infection, 5 mice were sacrificed from each vaccination and control group, and lungs were harvested to determine viral titer by a plaque assay and viral RNA levels by qRT-PCR.

#### SARS-CoV-2 plaque assay

SARS-CoV-2 lung titers were quantified by homogenizing harvested lungs in PBS (Quality Biological Inc.) using 1.0 mm glass beads (Sigma Aldrich) and a Beadruptor (Omni International Inc.). Homogenates were added to Vero E6 near confluent cultures and SARS-CoV-2 virus titers determined by counting plaque forming units (pfu) using a 6-point dilution curve.

### SARS-CoV-2 subgenomic RNA detection

Lung tissue was harvested and homogenized in TRIzol (Ambion). RNA was extracted per the manufacturer’s instructions using the Direct-zol RNA Miniprep Kit (Zymo Research). RNA was converted into cDNA (RevertAid Reverse Transcriptase) and used as template for qPCR (Applied Biosystems PowerUp SYBR Green Master Mix, Cat #A25742). The primers used were against the N gene (5⍰-TAATCAGACAAGGAACTGATTA-3⍰ (Forward) and 5⍰-CGAAGGTGTGACTTCCATG-3⍰ (Reverse)) on an Applied Biosystems QuantStudio 5 thermocycler. Data was analyzed in Prism 9 (Graphpad).

#### Anti-SARS-CoV-2 spike IgG by ELISA

An ELISA was used to determine anti-SARS-CoV-2 S IgG titers. Briefly, 96 well microtiter plates (ThermoFischer Scientific, Rochester, NY, USA) were coated with 1.0 µg mL^-1^ of SARS-CoV-2 spike protein. Plates were washed with PBS-T and blocked with TBS Startblock blocking buffer (ThermoFisher, Scientific). Mouse, baboon or human serum samples were serially diluted (10^−2^ to 10^−8^) and added to the blocked plates before incubation at room temperature for 2 hours. Following incubation, plates were washed with PBS-T and HRP-conjugated goat anti-mouse IgG or goat anti-human IgG (Southern Biotech, Birmingham, AL, USA) added for 1 hour. Plates were washed with PBS-T and 3,3’,5,5’-tetramethylbenzidine peroxidase substrate (TMB, T0440-IL, Sigma, St Louis, MO, USA) was added. Reactions were stopped with TMB stop solution (ScyTek Laboratories, Inc. Logan, UT). Plates were read at OD 450 nm with a SpectraMax®Plus plate reader (Molecular Devices, Sunnyvale, CA, USA) and data analyzed with SoftMax® (Molecular Devices, Corp.) software. EC_50_ values were calculated by 4-parameter fitting using SoftMax Pro 6.5.1 GxP software. Individual animal anti-SARS-CoV-2 rS IgG titers and group geometric mean titers (GMT) and 95% confidence interval (± 95% CI) were plotted GraphPad® Prism 7.05 software (Graphpad Software LLC).

#### hACE2 receptor blocking antibodies

Human ACE2 receptor blocking antibodies were determined by ELISA. Ninety-six well plates were coated with 1.0 μg mL^-1^ SARS-CoV-2 rS protein overnight at 4°C. After washing with PBS-T and blocking with StartingBlock (TBS) blocking buffer (ThermoFisher Scientific), serially diluted serum from groups of immunized mice or baboons were added to coated wells and incubated for 1 hour at room temperature. After washing, 30 ng mL^-1^ of histidine-tagged hACE2 (Sino Biologics, Beijing, CHN) was added to wells for 1 hour at room temperature. After washing, HRP-conjugated anti-histidine IgG (Southern Biotech, Birmingham, AL, USA) was added, followed by washing and addition of TMB substrate. Plates were read at OD 450 nm with a SpectraMax plus plate reader (Molecular Devices, Sunnyvale, CA, USA) and data analyzed with SoftMax Pro 6.5.1 GxP software. The % Inhibition for each dilution for each sample was calculated using the following equation in the SoftMax Pro program: 100-[(MeanResults/ControlValue@PositiveControl)*100].

Serum dilution versus % Inhibition plot was generated and curve fitting was performed by 4 parameter logistic (4PL) curve fitting to data. Serum antibody titer at 50% inhibition (IC_50_) of hACE2 to SARS-CoV-2 rS protein (rS-WU1 or rS-B.1.351) was determined in the SoftMax Pro program.

### SARS-CoV-2 Neutralization Titer by Plaque Reduction Neutralization Titer Assay (PRNT)

PRNTs were processed as described previously(*22*). Briefly, serum samples were diluted in DMEM (Quality Biological) at an initial 1:40 (baboon or mouse samples) or 1:100 (human samples) dilution with 1:2 serial dilutions for a total of 11 dilutions. A no-sera control was included on every plate. SARS-CoV-2 (WA-1, B1.1.7, or B1.351) was then added 1:1 to each dilution for a target of 50 PFU per plaque assay well and incubated at 37⍰C (5.0% CO_2_) for 1 hr. Sample titers where then determined by plaque assay and neutralization titers determined as compared to the non-treatment control. A 4-parameter logistic curve was fit to these neutralization data in PRISM (GraphPad, San Diego, CA) and the dilution required to neutralize 50% of the virus (PRNT50) was calculated based on that curve fit.

#### Surface and intracellular cytokine staining

For surface staining, murine splenocytes were first incubated with an anti-CD16/32 antibody to block the Fc receptor. To characterize T follicular helper cells (Tfh), splenocytes were incubated with the following antibodies or dye: BV650-conjugated anti-CD3, APC-H7-conjugated anti-CD4, FITC-conjugated anti-CD8, Percp-cy5.5-conjugated anti-CXCR5, APC-conjugated anti-PD-1, Alexa Fluor 700-conjugated anti-CD19, PE-conjugated anti-CD49b (BD Biosciences, San Jose, CA) and the yellow LIVE/DEAD^®^ dye (Life Technologies, NY). To stain germinal center (GC) B cells, splenocytes were labeled with FITC-conjugated anti-CD3, PerCP-Cy5.5-conjugated anti-B220, APC-conjugated anti-CD19, PE-cy7-conjugated anti-CD95, and BV421-conjugated anti-GL7 (BD Biosciences) and the yellow LIVE/DEAD^®^ dye (Life Technologies, NY).

For intracellular cytokine staining (ICCS) of murine splenocytes, cells were cultured in a 96-well U-bottom plate at 2 × 10^6^ cells per well. The cells were stimulated with rS-WU1 or rS-B.1.351 spike protein. The plate was incubated 6 h at 37°C in the presence of BD GolgiPlug™ and BD GolgiStop™ (BD Biosciences) for the last 4 h of incubation. Cells were labeled with murine antibodies against CD3 (BV650), CD4 (APC-H7), CD8 (FITC), CD44 (Alexa Fluor 700), and CD62L (PE) (BD Pharmingen, CA) and the yellow LIVE/DEAD^®^ dye. After fixation with Cytofix/Cytoperm (BD Biosciences), cells were incubated with PerCP-Cy5.5-conjugated anti-IFN-γ, BV421-conjugated anti-IL-2, PE-Cy7-conjugated anti-TNF-α, and APC-conjugated anti-IL-4 (BD Biosciences). All stained samples were acquired using a LSR-Fortessa or a FACSymphony flow cytometer (Becton Dickinson, San Jose, CA) and the data were analyzed with FlowJo software version Xv10 (Tree Star Inc., Ashland, OR).

For ICS of baboon PBMCs, PBMCs collected at the timepoints listed in Figure 5A were stimulated as described above with rS-WU1 or rS-B.1.351. Cells were labelled with human/NHP antibodies BV650-conjugated anti-CD3, APC-H7-conjugated anti-CD4, FITC-conjugated anti-CD8, BV421-conjugated anti-IL-2, PerCP-Cy5.5-conjugated anti-IFN-γ, PE-Cy7-conjugated anti-TNF-α, APC-conjugated anti-IL-5, BV711-conjugated anti-IL-13 (BD Biosciences), and the yellow LIVE/DEAD^®^ dye.

### ELISpot Assay

Murine IFN-γ and IL-5 ELISpot assays were performed following the manufacturer’s procedures for mouse IFN-γ and IL-5 ELISpot kits (3321-2H and 3321-2A, Mabtech, Cincinnati, OH). Briefly, 4 × 10^5^ splenocytes in a volume of 200⍰µL were stimulated with rS-WU1 or rS-B.1.351 in plates that were pre-coated with anti-IFN-γ or anti-IL-5 antibodies. Each stimulation condition was carried out in triplicate. Assay plates were incubated 24-48 h at 37⍰°C in a 5% CO_2_ incubator and developed using BD ELISpot AEC substrate set (BD Biosciences, San Diego, CA). Spots were counted and analyzed using an ELISpot reader and ImmunoSpot®software v6 (Cellular Technology, Ltd., Shaker Heights, OH). The number of IFN-γ- or IL-5-secreting cells was obtained by subtracting the background number in the medium controls. Data shown in the graph are the average of triplicate wells.

Similarly, baboon IFN-γ and IL-4 assays were carried out using NHP IFN-γ and Human IL-4 assay kits from Mabtech. Assays were performed in triplicate.

#### Statistical analysis

Statistical analyses were performed with GraphPad Prism 8.0 software (La Jolla, CA). Serum antibody titers were plotted for individual animals and the geometric mean titer (GMT) and 95% confidence interval (95% CI) or the means ± SEM as indicated in the figure. Ordinary one-way ANOVA with Tukey’s multiple comparisons post-hoc test was performed on log_10_-transformed data to evaluate statistical significance of differences among groups. P-values ≤0.05 were considered as statistically significant.

## Data availability

The authors declare that all data supporting the findings of the study are available in the article and the Supplementary Information file.

## Acknowledgements

Authors would like to acknowledge Joyce Plested and Shane Cloney-Clark from Novavax Clinical Immunology Department for providing human serum samples for microneutralization assays. Novavax funded the work at UMSOM.

## Supplemental Figs

**Figure S1.**
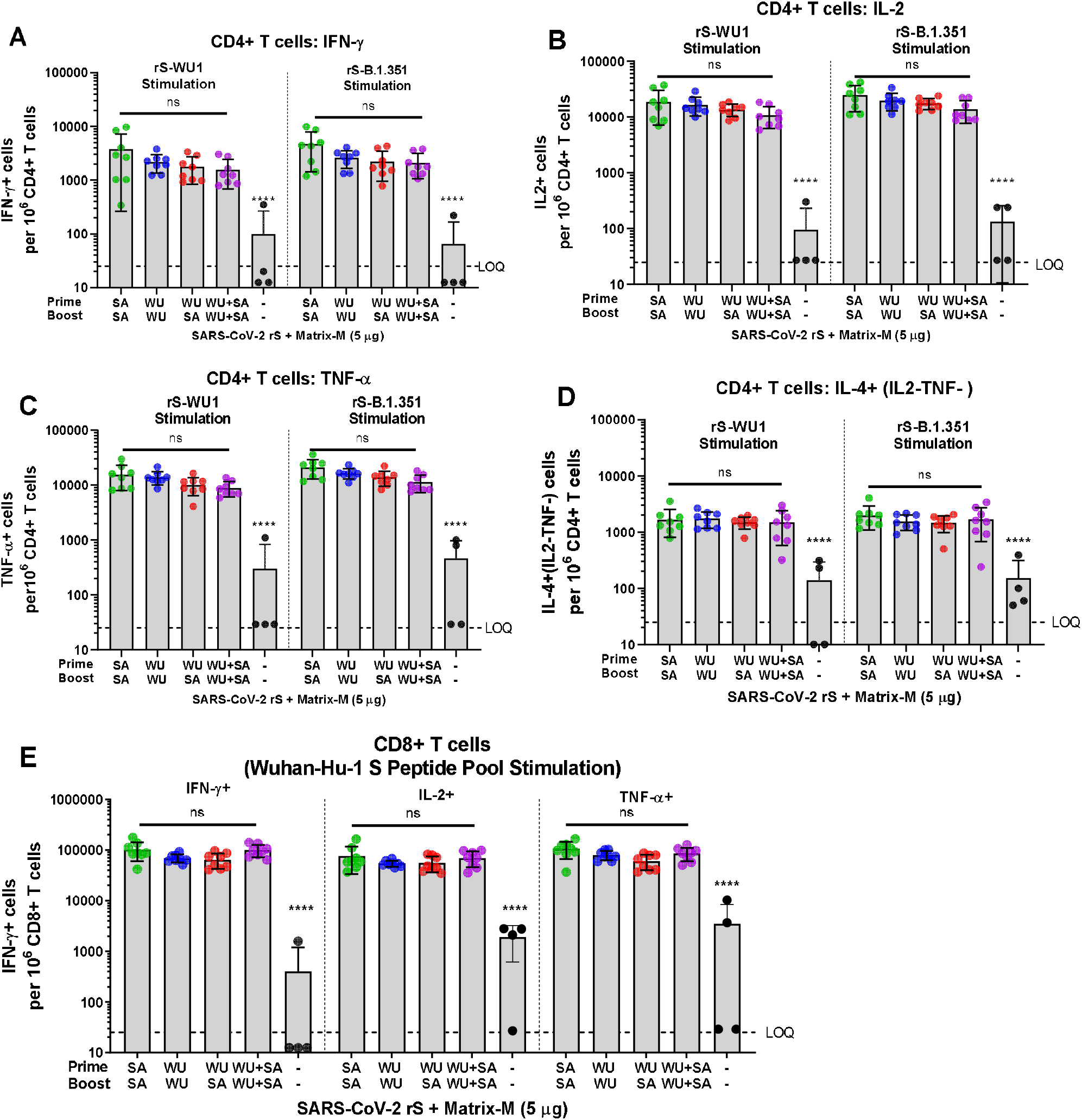
Individual Cytokine responses in Mice. Groups of mice (n = 8/group) were immunized in a prime/boost regimen on Days 0 and 21 with various combinations of SARS-CoV-2 rS based on B.1.351 (SA) or Wuhan-Hu-1 (WU). Mice were either primed and boosted with rS-B.1.351, primed and boosted with rS-WU1, primed with rS-WU1 and boosted with rS-B.1.351, or primed and boosted with bivalent rS-WU1+rS-B.1.351. Antigen doses were 1 µg rS for each monovalent immunization, or 1 µg rS for each construct upon bivalent immunization (2 µg rS total). All antigen doses were administered with 5 µg Matrix-M1 adjuvant. A control group received formulation buffer (Placebo, n = 5). Spleens were harvested on Day 28 for cell collection. **A-D**. Isolated splenocytes were stimulated with either rS-WU1 or rS-B.1.351, then subjected to intracellular cytokine staining to determine whether CD4+ T cells were positive for IFN-γ (**A**), IL-2 (**B**), TNF-α (**C**), or IL-4 (**D**). **E**. To examine CD8+ T cell responses, cells were stimulated with a peptide pool corresponding to the entire Wuhan-Hu-1 spike protein sequence, then subjected to ICS for IFN-γ, IL-2, and TNF-α.

**Figure S2.**
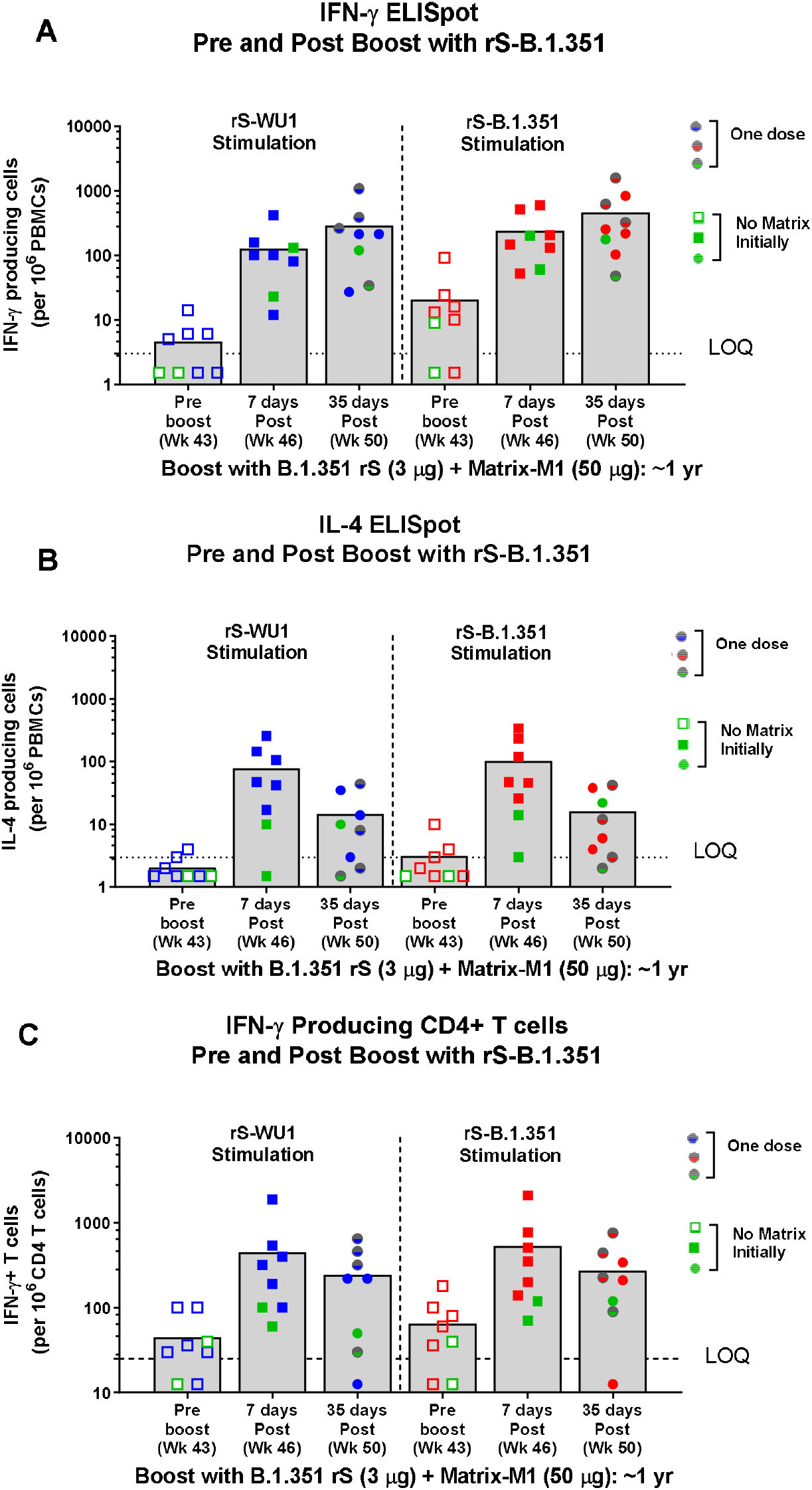
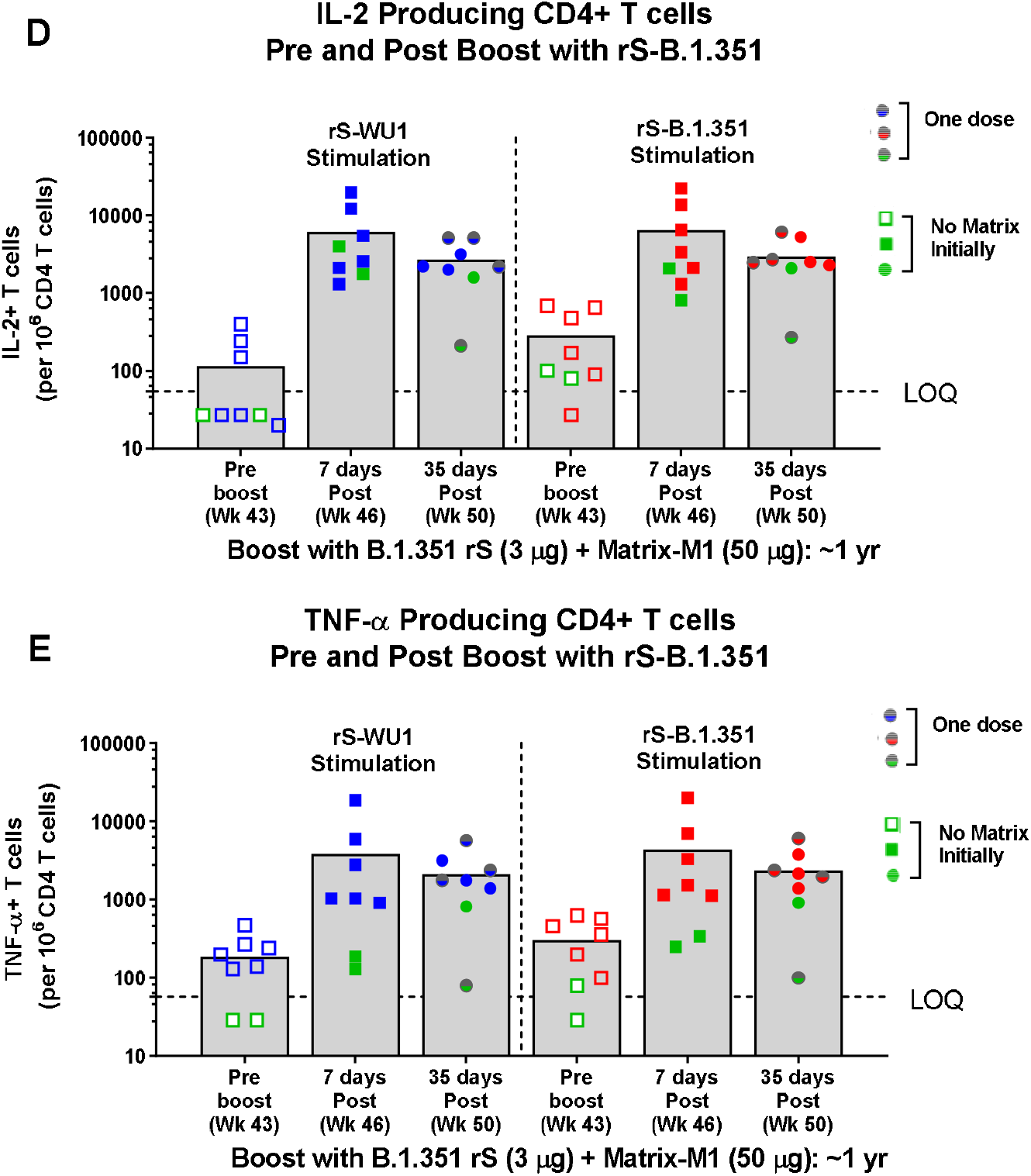
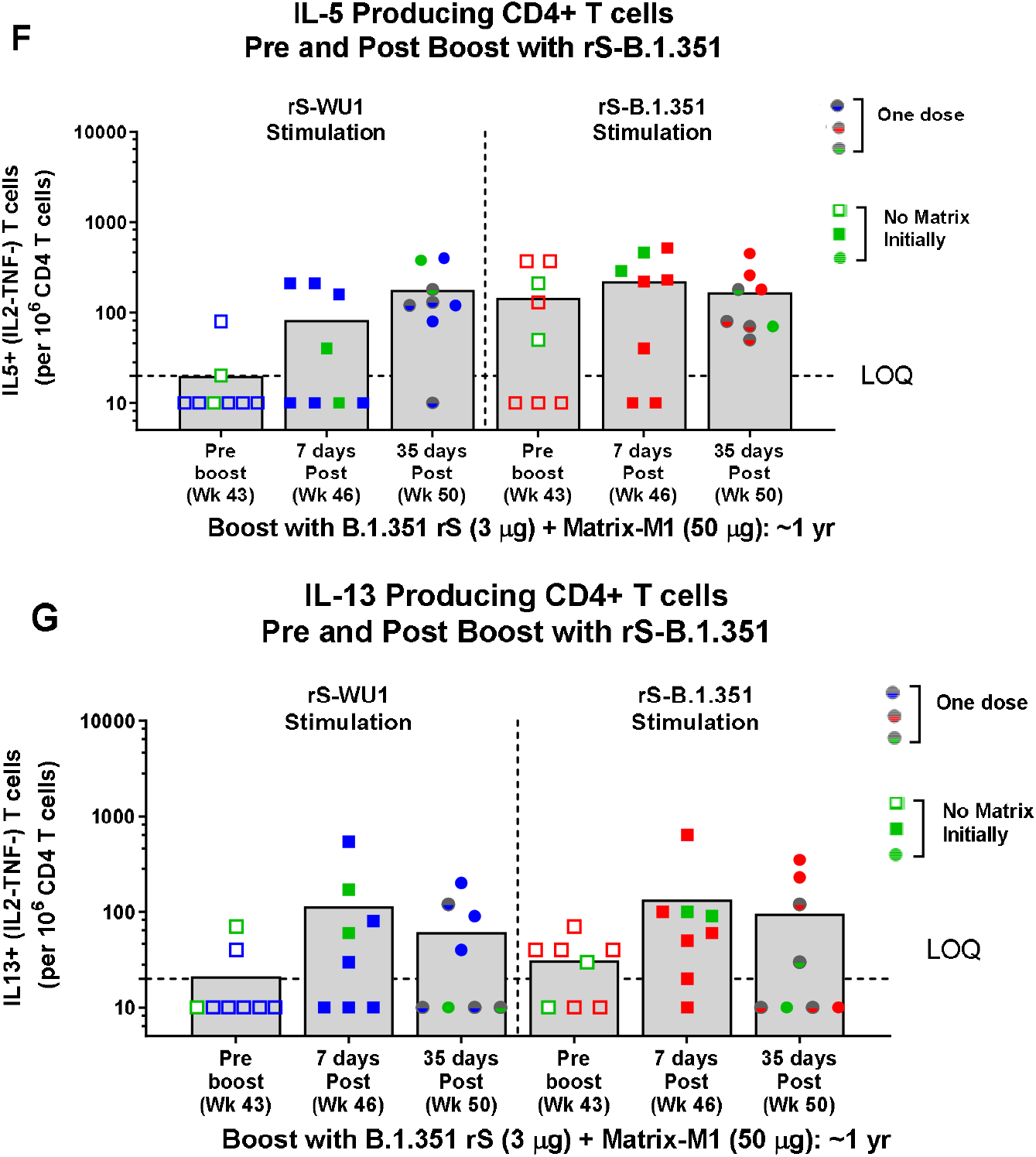
Individual Cytokine responses to NVX-CoV2438 boost in Baboons. Individual Cytokine Responses to rS-B.1.351 boost in Baboons. A small cohort of baboons (n = 2-3/group) was immunized with 1 µg, 5 µg, or 25 µg rS-WU1 with 50 µg Matrix-M1 adjuvant or unadjuvanted 25 µg rS-WU1 on Day 0 and 21 (Week 0 and 3, respectively). Approximately 1 year later, all animals were boosted with one or two doses of 3 µg rS-B.1.351 with 50 µg Matrix-M1 adjuvant on Day 318 and 339 (Weeks 45 and 48, respectively). PBMCs collected pre-boost (Day 303; Week 43), 7 days after the first rS-B.1.351 boost (Day 325; Week 46), and 35 days after the first rS-B.1.351 boost (Day 353; Week 50). PBMCs were stimulated with rS-WU1 or rS-B.1.351 and subjected to ELISpot assay to measure **(A)** IFN-γ producing cells as a Th1 cytokine and **(B)** IL-4 producing cells as a Th2 cytokine. CD4+ T cells were also stimulated with rS-WU1 or rS-B.1.351, then subjected to ICS to measure cells producing IFN-γ **(C)**, IL-2 **(D)**, TNF-α **(E)**, IL-5 **(F)**, and IL-13 **(G)**.

